# *Hypericum triquetrifolium* Essential Oil: Chemical Composition, Hierarchical Cluster Analysis, and Antibacterial Activity

**DOI:** 10.1101/2025.03.12.642897

**Authors:** Tamam El-Elimat, Haya S. El-Qaderi, Mohammad Hudaib, Ahmed H. Al Sharie, Myassar O. Alekish, Mohammad Al-Gharaibeh, Wael M. Hananeh

**Author notes:** Corresponding author. *E-mail address:* (T. El-Elimat).

## Abstract

Plant species belonging to the genus *Hypericum* are well known for their profound pharmacological activities. This study evaluated the *in vitro* antibacterial activity and the chemical composition of the essential oil of *Hypericum triquetrifolium* Turra (Hypericaceae) that was collected from the Northern part of Jordan. The aerial parts of *H. triquetrifolium* were subjected to hydrodistillation to obtain a yellowish essential oil with a yield of 0.023% (w/w of dried material). The chemical composition of the oil was explored using gas chromatography–mass spectrometry (GC-MS). Forty-five compounds accounting for 99.5% of the oil composition were identified. The main classes identified were monoterpenes with 52.3% (49.6% as monoterpene hydrocarbons and 2.7% as oxygenated monoterpenes), aliphatic hydrocarbons 35.4%, and sesquiterpenes 11.8% (10.9% as sesquiterpene hydrocarbons and 0.9% as oxygenated sesquiterpenes). The oil was predominated with *α*-pinene (40.0%), which was identified as a chemotype for Jordanian *H. triquetrifolium*. Other major constituents included n-nonanal (10.1%), nonane (9.0%), 4*Z*-hepten-1-ol (5.8%), 4-methyl-pentanol (5.1%), and *β*-pinene (4.6%). The antimicrobial activity of the oil was evaluated using a micro broth dilution assay against a set of Gram-positive and Gram-negative pathogens. The oil exhibited antimicrobial activity against the tested strains with minimum bactericidal concentration (MBC)/minimum inhibitory concentration (MIC) values of 12.5/6.25, 6.25/3.125, 6.25/3.125 and 3.125/1.56 *μ*g/*μ*L, respectively. Finally, Hierarchical Cluster Analysis of the essential oil composition of *H. triquetrifolium* from different geographical locations in the world has identified the oil from Jordan to be more similar to those obtained from Italy, Greece, and Crete.

## 1. Introduction

*Hypericum triquetrifolium* Turra (*H. crispum* L.) is a flowering plant belongs to the family Hypericaceae, a family of 8 genera and more than 600 species that are distributed all over the world [1]. The genus *Hypericum* is the largest among the eight genera constituting more than 500 species, including small trees, shrubs, and herbs [1, 2]. The genus has a worldwide distribution flourishing mainly in warm-temperate areas [2, 3]. One of the world-renowned top-selling medicinal plants that belongs to the genus *Hypericum* is St. John’s wort (*Hypericum perforatum*). It is widely used as an alternative medicine for treatment of mild to moderate depression [4]. *Hypericum* is well represented in Jordan; Al-Eisawi in his book “*Flora of Jordan*” has reported eight species growing in the wild, namely: *H. geslini* Coss. & Karl, *H. languginosum* Lam., *H. olivieri* (Spach) Boiss., *H. perforatum* Post, *H. serpyllifolium* Lam., *H. sinaicum* Hochst. Ex Boiss., *H. thymifolium* Banks & Sol., and *H. triquetrifolium* Turra [5].

*H. triquetrifolium* is a wild growing weed in Jordan that is native to Northern Africa, Western Asia, and Southeastern and Southwestern Europe [5, 6]. It is commonly referred to as wavy-leaf St. John’s wort, curled leaf St. John’s wort, tangled Hypericum; Dirnah (Lebanon), and locally known in Jordan as “Roja” or “Aran”. *H. triquetrifolium*, which is found flowering from May to August, is a perennial herb, which grows up to 50 cm high with a dense tangle of thin branches, smooth but dotted with small black glands. The leaves are opposite and simple, 5-15 mm long, with wavy ends. Flowers are stalked in groups of 2-5 at the branches end, with 5 free yellow petals [7].

*H. triquetrifolium* is reported to have a wide spectrum of pharmacological activities, including cytotoxic, anti-inflammatory, antinociceptive, antibacterial, and antioxidant activities [8, 9, 10, 11, 12, 13, 14, 15], which could be attributed to its content of secondary metabolites, such as phloroglucinols, naphthodianthrones, and flavonoids [8, 13, 14, 16, 17]. In traditional medicine of Jordan, the plant is reported to have sedative, astringent, and antispasmodic effects and was used for intestine and bile disorders [18].

Most of the pharmacological studies of the *Hypericum* species were directed toward the crude extracts. However, the essential oils extracted by steam distillation had received little attention in terms of clinical data to proof its safety and efficacy [19]. Plants belong to the genus *Hypericum* are known to be poor producers of essential oils with a percentage yield less than 1% w/w [19].

The chemical composition of the essential oil of *H. triquetrifolium* collected from different locations in the Mediterranean region (Turkey, Tunisia, Iran, and Iraq) was studied by gas chromatography (GC) and by gas chromatography-mass spectrometry (GC-MS) [20, 21, 22, 23, 24, 25]. Germacrene-D, *β*-caryophyllene, *α*-pinene, and caryophyllene oxide were found to be the predominant constituents [20, 21, 22, 23, 24, 25]. As the chemical composition of *H. triquetrifolium* essential oils showed variability quantitatively and qualitatively due to many intrinsic and extrinsic environmental factors, including geographical locations [6], and as Jordan has a unique position in the heart of the Middle East at the junction of three continents (Asia, Africa, and Europe), which bestows the country with rich biodiversity of wild plants [26]. Hence, we sought to explore the chemical composition and the antimicrobial activity of *H. triquetrifolium* essential oil growing wild in the Northern part of Jordan.

## 2. Results and discussion

### 2.2 Gas chromatography-mass spectrometry (GC-MS) analysis of the essential oil of H. triquetrifolium

Hydro-distillation of the aerial parts of *H. triquetrifolium* grown in Jordan furnished a yellowish essential oil with a yield of 0.023% (w/w of dried material). The chemical composition of the obtained essential oil was determined by GC-MS (Table 1 and Figure 1). Comparison of the GC retention (Kovat’s) indices calculated using n-alkane hydrocarbon mixture with those reported in the literature [27] enabled the identification of 99.5% of the oil composition. A total of 45 compounds were identified belonging to three major classes, namely monoterpenes with 52.3% (49.6% as monoterpene hydrocarbons and 2.7% as oxygenated monoterpenes), aliphatic hydrocarbons 35.4%, and sesquiterpenes 11.8% (10.9% as sesquiterpene hydrocarbons and 0.9% as oxygenated sesquiterpenes). The rest of the oil components (0.5%) were either traces or unidentified. The major compounds identified were *α*-pinene (40.0%), n-nonanal (10.1%), nonane (9.0%), 4*Z*-hepten-1-ol (5.8%), 4-methyl-pentanol (5.1%), *β*-pinene (4.6%), n-octanol (3.5%), limonene (2.1%), and α-himachalene (2.0%).

**Table 1.** Chemical composition of the essential oil obtained by hydrodistillation of the aerial parts of *H. triquetrifolium*.

| # | LI | CI | Compound | %content | Method of Identification |
| --- | --- | --- | --- | --- | --- |
| 1 | 838 | 838 | 4-Methyl-pentanol | 5.1 | RI-MS |
| 2 | 900 | 901 | Nonane | 9.0 | RI-MS |
| 3 | 901 | 904 | Ethyl pentanoate | tr | RI-MS |
| 4 | 939 | 932 | α-Pinene | 40.0 | RI-MS |
| 5 | 967 | 968 | 4Z-hepten-1-ol | 5.8 | RI-MS |
| 6 | 979 | 979 | β-Pinene | 4.6 | RI-MS |
| 7 | 986 | 984 | 6-Methyl-5-hepten-2-one | tr | RI-MS |
| 8 | 991 | 989 | β-Myrcene | 0.8 | RI-MS |
| 9 | 1000 | 1001 | Decane | 0.3 | RI-MS |
| 10 | 1025 | 1025 | p-Cymene | 1.6 | RI-MS |
| 11 | 1029 | 1029 | Limonene | 2.1 | RI-MS |
| 12 | 1060 | 1058 | γ-Terpinene | 0.7 | RI-MS |
| 13 | 1068 | 1064 | n-Octanol | 3.5 | RI-MS |
| 14 | 1089 | 1086 | Terpinolene | 0.4 | RI-MS |
| 15 | 1091 | 1090 | p-Cymenene | 0.2 | RI-MS |
| 16 | 1101 | 1101 | n-Nonanal | 10.1 | RI-MS |
| 17 | 1126 | 1127 | α-Campholenal | 0.3 | RI-MS |
| 18 | 1131 | 1132 | Octyl formate | 0.3 | RI-MS |
| 19 |  | 1136 | Unknown | 0.5 | --- |
| 20 | 1139 | 1142 | *trans*-Pinocarveol | 0.6 | RI-MS |
| 21 | 1169 | 1172 | 3-Thujanol | 0.7 | RI-MS |
| 22 | 1189 | 1195 | α-Terpineol | 0.6 | RI-MS |
| 23 | 1199 | 1203 | γ-Terpineol | 0.3 | RI-MS |
| 24 | 1217 | 1219 | *trans*-Carveol | 0.2 | RI-MS |
| 25 | 1300 | 1300 | n-Tridecane | 0.3 | RI-MS |
| 26 | 1375 | 1367 | α-Ylangene | 0.2 | RI-MS |
| 27 | 1377 | 1373 | α-Copaene | 0.3 | RI-MS |
| 28 | 1421 | 1414 | β-Cedrene | 0.7 | RI-MS |
| 29 | 1419 | 1418 | β-Caryophyllene | 0.4 | RI-MS |
| 30 | 1441 | 1436 | Aromadendrene | 0.1 | RI-MS |
| 31 | 1447 | 1443 | Seychellene | 0.8 | RI-MS |
| 32 | **1451** | **1448** | **α-Himachalene** | **2.0** | **RI-MS** |
| 33 | 1460 | 1454 | Allo-aromadendrene | 0.5 | RI-MS |
| 34 | 1480 | 1473 | γ-Muurolene | 1.2 | RI-MS |
| 35 | 1483 | 1475 | γ-Himachalene | 1.2 | RI-MS |
| 36 | 1481 | 1480 | α-Curcumene | 0.2 | RI-MS |
| 37 | 1496 | 1490 | γ-Amorphene | 0.3 | RI-MS |
| 38 | 1505 | 1498 | β-Himachalene | 1.4 | RI-MS |
| 39 | 1510 | 1506 | Tridecanal (impure) | 0.2 | RI-MS |
| 40 | 1514 | 1511 | γ-Cadinene | 0.7 | RI-MS |
| 41 | 1523 | 1516 | δ-Cadinene | 1.0 | RI-MS |
| 42 | 1529 | 1520 | *trans*-Calamenene | 0.3 | RI-MS |
| 43 | 1535 | 1531 | *trans*-Cadina-1(2),4-diene | 0.1 | RI-MS |
| 44 | 1539 | 1535 | α-Cadinene | 0.1 | RI-MS |
| 45 | 1546 | 1539 | α-Calacorene | 0.1 | RI-MS |
| 46 | 1616 | 1610 | β-Himachalene oxide | 0.2 | RI-MS |
| **Aliphatic hydrocarbons** |  |  |  | **35.4** |  |
| **Monoterpenes** |  |  |  | **52.3** |  |
| Monoterpene hydrocarbons |  |  |  | 49.6 |  |
| Oxygenated monoterpenes |  |  |  | 2.7% |  |
| **Sesquiterpenes** |  |  |  | **11.8** |  |
| Sesquiterpene hydrocarbons |  |  |  | 10.9 |  |
| Oxygenated sesquiterpenes |  |  |  | 0.9 |  |
| **Traces and Unknown** |  |  |  | **0.5** |  |

Rt: Retention time, CI: calculated arithmetic Kovat's index on DB-5 column, LI: arithmetic Kovat's index, based on DB-5 column, as reported in literature (Adams, 2007); compounds in bold are the major oil components (%content $\geq 2\%$ ); tr: traces amount $< 0.1\%$ ; Unknown oxygenated hydrocarbon (CI 1136).

**Figure 1.**
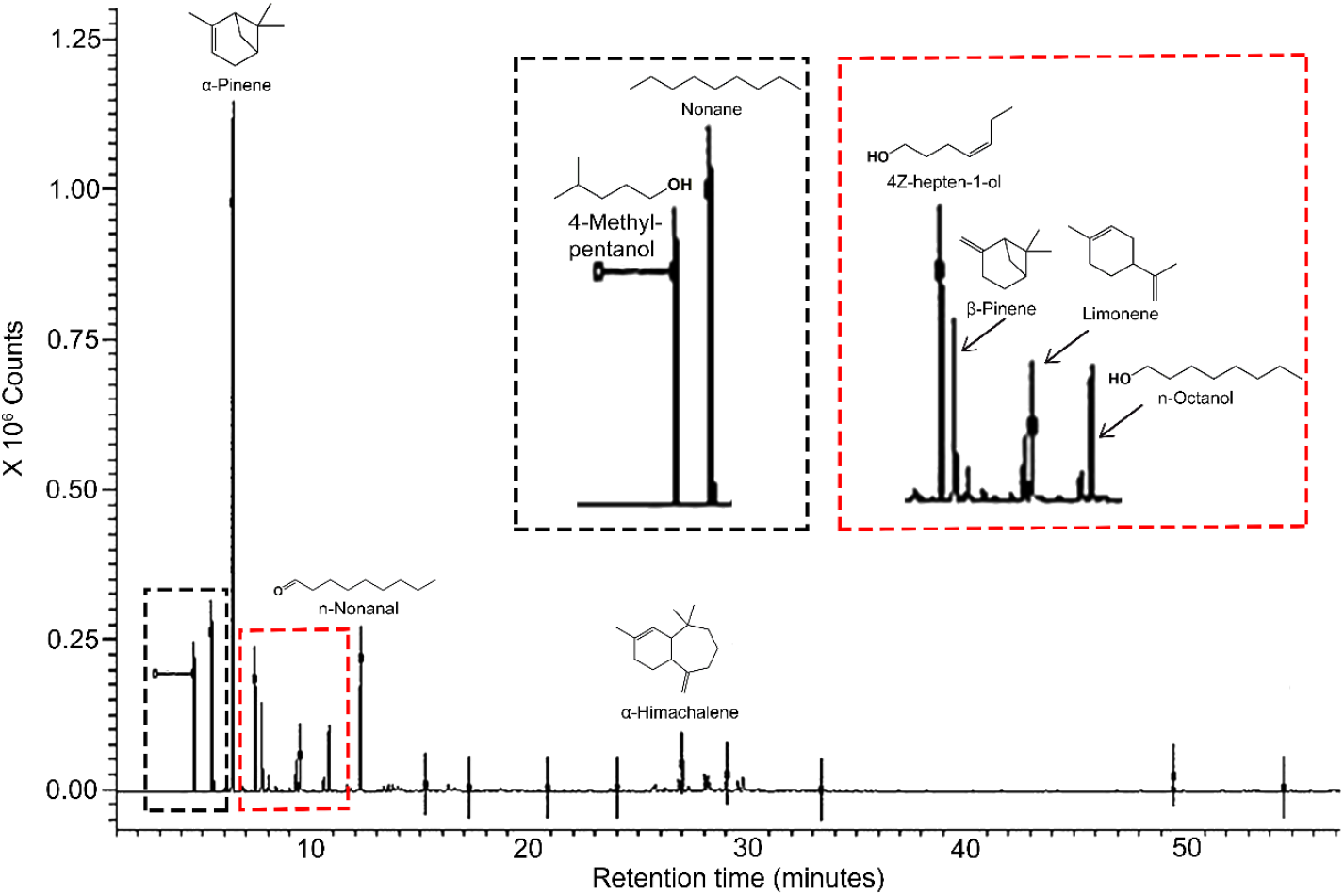
GC-MS chromatogram of the essential oil hydrodistilled from *H. triquetrifolium* aerial parts.

The principal monoterpene hydrocarbon that dominated the oil composition was *α*-pinene (40%), which could be identified as a chemotype of Jordanian *H. triquetrifolium*. However, oxygenated monoterpenes were represented by six compounds with a concentration of less than 1% for each. On the other hand, the hydrocarbons are represented by ten compounds with n-nonanal and nonane were the major ones. Sesquiterpene hydrocarbon class of compounds was the most diverse among other classes with 18 compounds have been identified, including *α*-himachalene, *β*-himachalene, *γ*-himachalene, and *δ*-cadinene. Oxygenated sesquiterpenes were the least among other classes with only two compounds, both constituting less than 1%.

The chemical composition of the essential oil of *H. triquetrifolium* collected from different locations was studied previously using GC and GC-MS. One hundred and nine compounds were identified from the essential oil of the aerial part of Tunisian *H. triquetrifolium* consisting of 92.2% of total detected constituents [20]. Main constituents were sesquiterpenes hydrocarbons (59.37%); main compounds in this class were *α*-humulene, *cis*-calamenene, *δ*-cadinene, bicyclogermacrene, eremophilene, *β*-caryophyllene, and (*E*)-*γ*-bisabolene. Monoterpene hydrocarbons (12.19%) with the main compound was *α*-pinene (10.33%). The oxygenated sesquiterpenes (9.33%) was mainly consisted of caryophyllene oxide (1.38%); while the oxygenated monoterpenes was weakly represented (4.62%) and contains compounds with low percentages (<1%) [20]. Another study on Tunisian oil identified one hundered seventy four compounds with the sesquiterpenes being the most abundant chemical compounds [23]. The major components were *β*-caryophyllene, *α*-humulene, *γ*-muurolene, germacrene D, *β*-selinene, valencene, *α*-muurolene, *trans*-*γ*-cadinene, and *δ*-cadinene [23].

Fifty compounds were identified in an Iranian essential oil consisting mainly of sesquiterpenes (85%) [24]. The major components were germacrene-D, *β*-caryophyllene, *δ*-cadinene, *trans*-*β*-farnesene, *α*-humulene, *β*-selinene, *γ*-cadinene and *trans*-phytol [24].

In Turkey, analysis of the essential oil of *H. triquetrifolium* showed that the monoterpene concentrations were higher than that of sesquiterpenes [25]. Forty five compounds were identified; 1-hexanal (18.8%), 3-methylnonane (12.5%), and *α*-pinene (12.3%) were the main compounds [25]. A summary of previously isolated and chemically analyzed essential oils from *H. triquetrifolium* in the Mediterranean region is summarized in Table 2.

**Table 2.**
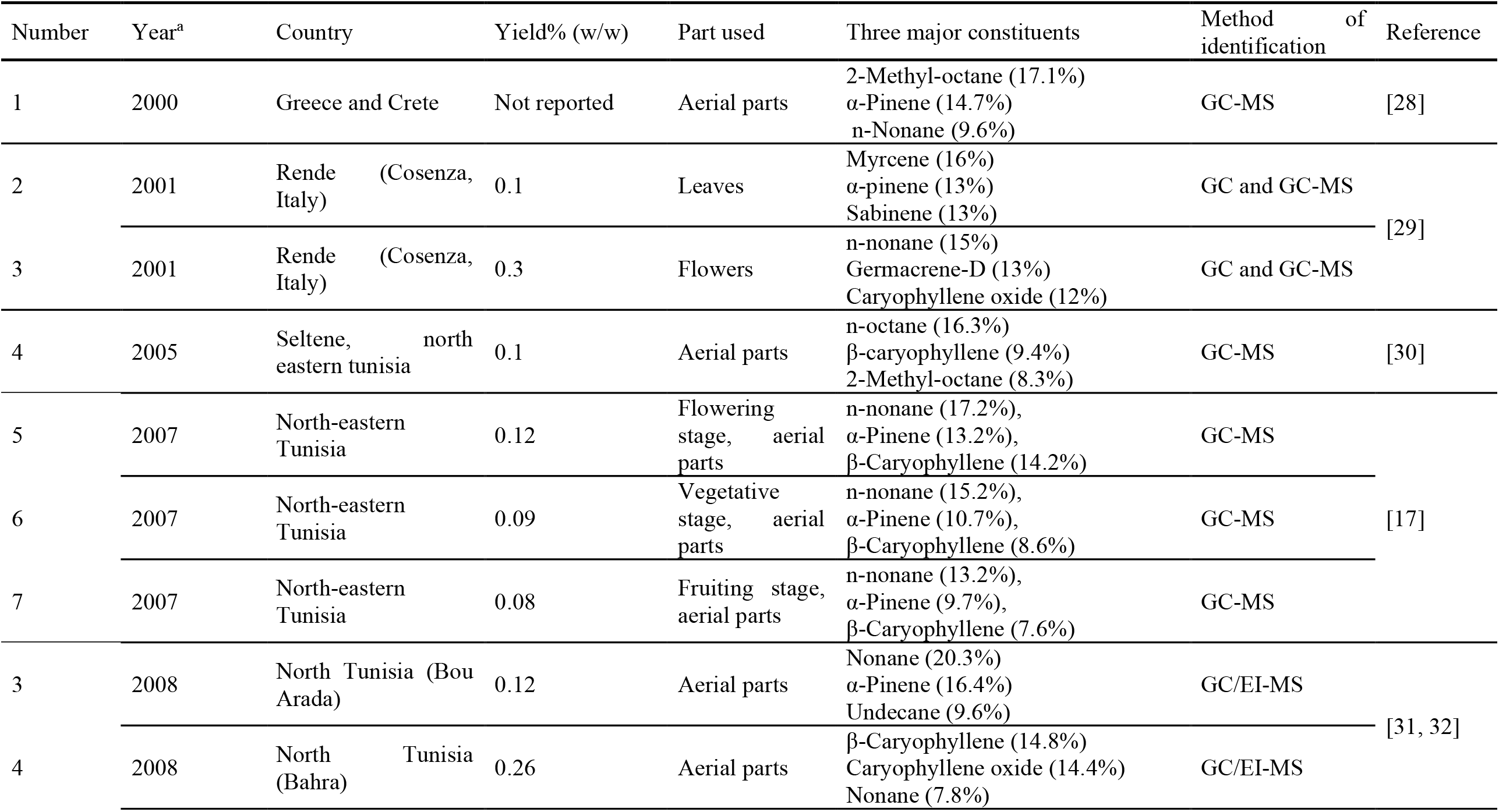
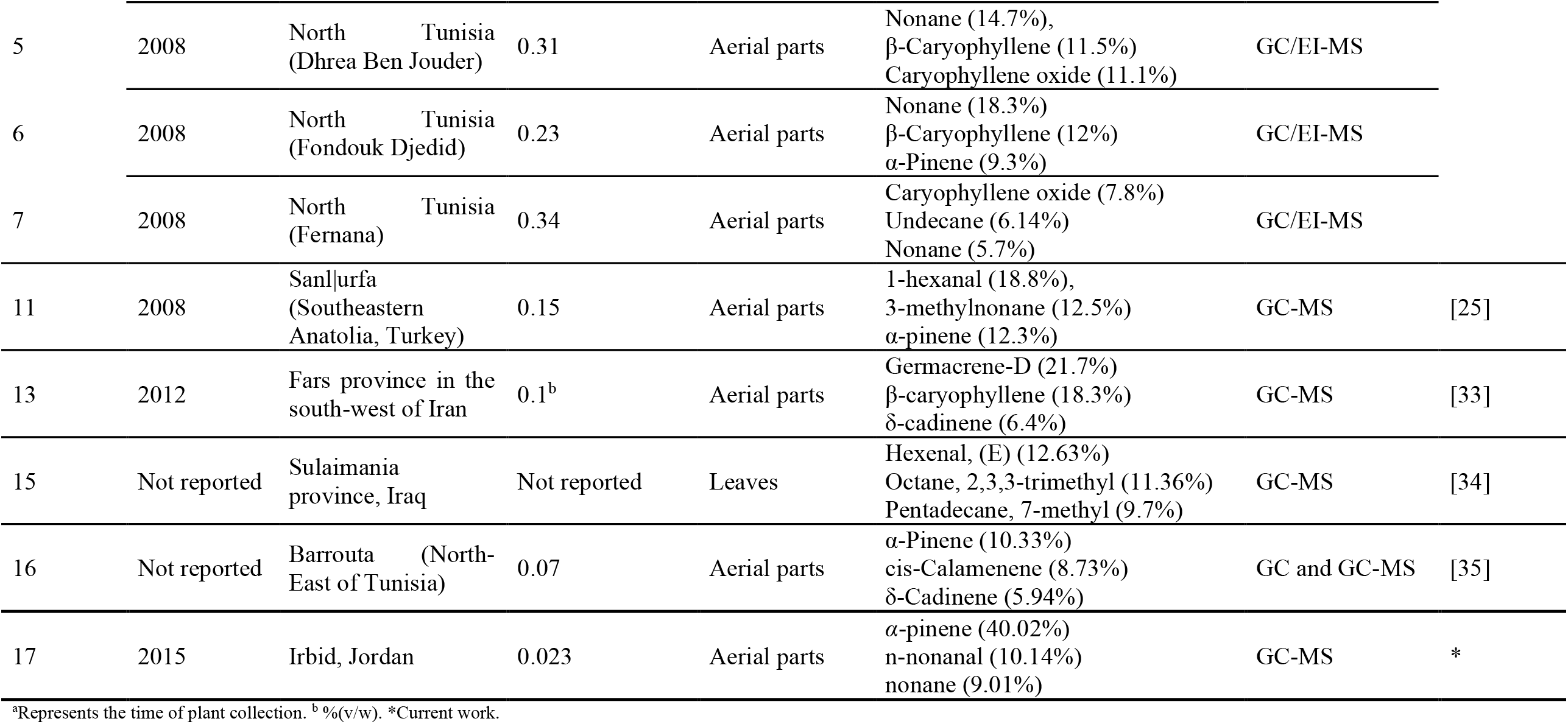
Systematic summary of previously isolated and chemically analyzed essential oils from *H. triquetrifolium* in the Mediterranean region.

The current study was consistent with the Iranian study in the fact that the essential oil of *H. triquetrifolium* consists mainly of monoterpenes even though the chemical composition is different. However, the oil composition is different from those obtained from Tunisia and Turkey where the oils were dominated by sesquiterpenes.

### 2.2 Hierarchical cluster analysis

To get a better chemotype comparison between oils obtained from different geographical locations in the world, the essential oils compositions were analysed by Hierarchical Cluster Analysis (HCA) as shown in Figure 2. Samples extracted using hydrodistillation were only involved in this study. A total of 16 samples and 312 compounds were identified, which were clustered into three separate clusters, namely C1, C2, and C3. The first cluster (C1) is obtained mainly by the essential oil extracted in five geographical locations in Tunisia (Figure 2). The second cluster (C2) is generated by clustering three oil samples extracted from the same location at different stages in Tunisia, while the other oil samples are also two Tunisian ones (Figure 2). The essential oil chemical profile from Jordan (this work) was clustered firstly with the samples extracted from Italy, Greece, and Crete, followed by Turkish and Iranian samples (Figure 2) to form cluster number 3 (C3). These data were indicative that the essential oil from Jordan is chemically more similar to those from Italy, Greece, and Crete, followed by Turkey and Iran. However, it is different from that of Tunisia. Intestinally, the Tunisian oil is standing by itself away from other oils analysed in this study.

**Figure 2.**
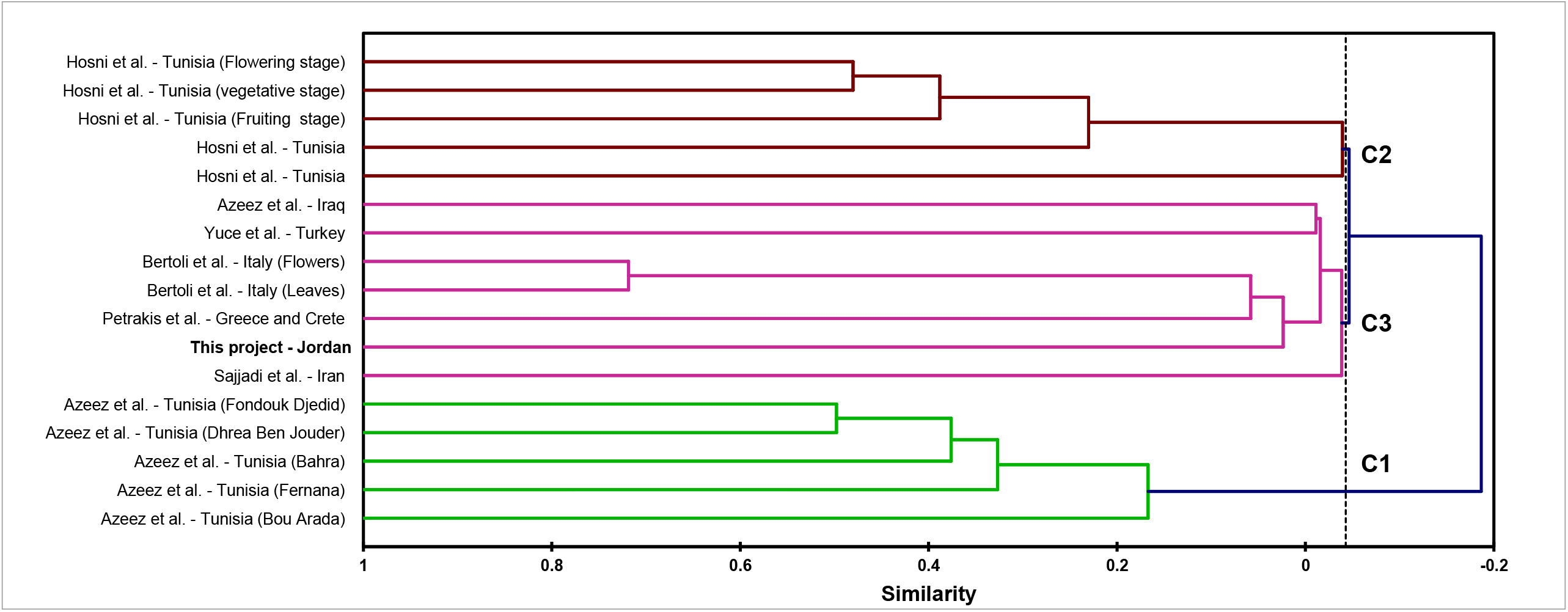
Dendrogram generated by agglomerative hierarchical clustering of 17 essential oil chemical profiles of *H. triquetrifolium*.

### 2.3 Antibacterial activity of H. triquetrifolium essential oil

The antimicrobial activity of *H. triquetrifolium* essential oil was evaluated using disk diffusion and micro broth dilution assays. The disc diffusion test results were not reported here, as there was no clear zone of inhibition around discs, since the oil did not diffuse well through the agar. Therefore, the results of the micro broth dilution assay using the 96-well plate only were reported in this paper. The tested oil shows MBC/MIC values of 12.5/6.25, 6.25/3.125, 6.25/3.125 and 3.125/1.56 *μ*g/*μ*L against *E. coli, S. aureus, S. chromogenes*, and *B. subtilis*, respectively (Table 3). This effect could be explained, in part, by the high concentrations of *α*- and *β*-pinene, which possess potent antimicrobial activity as previously described [36]. The essential oil of *H. triquetrifolium* was previously studied for its antibacterial activity against eight bacterial strains [21]. Among *B. cereus, B. brevis, S. pyogenes, P. aeruginosa, E. coli*, and *S. aureus*; the essential oil was very active against *B. cereus, P. aeruginosa, E. coli*, and *S. aureus* at 80 *μ*g/paper disc. The inhibition zones 20, 20, 20 and 16, respectively were found to be more than the inhibition zones of the standard antibiotic (ampicillin/slubactam 10/10 *μ*g/paper disc), which was reported as 12, 14, 10 and 12, respectively [21]. For the other tested strains; the oil showed antibacterial activity, but with less or similar inhibition zone to the standard antibiotic [21]. Rouis, Z., et al. (2013) have studied the antibacterial activity for *H. triquetrifolium* essential oil from different regions in Tunisia against *B. cereus, E. coli, Vibrio alginolyticus, V. cholera, P. aeruginosa, Salmonella typhimurium, Aeromonas hydrophila, Enterococcus faecalis, S. aureus*, and *S. epidermidis* [22].

**Table 3.** Antibacterial activity of *H. triquetrifolium* essential oil.

| Bacterial strain | MBC ($\mu\text{g}/\mu\text{L}$) | MIC ($\mu\text{g}/\mu\text{L}$) |
| --- | --- | --- |
| *Staphylococcus aureus* | 6.25 | 3.125 |
| *Staphylococcus chromogenes* | 6.25 | 3.125 |
| *Escherichia coli* | 3.125 | 1.56 |
| *Bacillus subtilis* | 12.5 | 6.25 |

MIC: Minimum Inhibitory Concentration; MBC: Minimum Bactericidal Concentration.

## 3. Experimental

### 3.1 Plant material

The aerial parts of *H. triquetrifolium* were collected during flowering stage in July 2015 from different populations throughout the campus of Jordan University of Science and Technology (JUST), Irbid, Jordan. The collected plant material was identified by Dr. Mohammad Al-Gharaibeh, Plant Taxonomist, Faculty of Agriculture, JUST. A voucher specimen (PHS-120) was deposited at the herbarium of the Faculty of Pharmacy, JUST. The plant material was air-dried in shadow away from direct sunlight in a well-ventilated area. The dried plant material was ground to a powder using an electrical laboratory mill, stored at room temperature (24 ± 4 °C) protected from light until required for analysis.

### 3.2 Preparation of H. triquetrifolium essential oil

Around 250 g of finely ground air-dried aerial parts of *H. triquetrifolium* were subjected to hydrodistillation for 3 h using a Clevenger-type apparatus according to the British Pharmacopoeia [37]. The floral water (condensate) was subjected to a liquid-liquid extraction with of dichloromethane and transferred into a separatory funnel; extraction was repeated 3 times and the organic layer was drawn off and evaporated to dryness using a rotary evaporator under reduced pressure at 40 °C. The oil obtained [0.023% (w/w of dry weight)] was dried over anhydrous sodium sulfate and then stored at 4 °C in amber air-tight sealed vials until required for analysis.

### 3.3 Gas chromatography-mass spectrometry (GC-MS) analysis of the essential oil of H. triquetrifolium

Quantitative and qualitative analyses of the essential oil were carried out using GC-MS. GC-MS was performed by a Varian Chrompack CP-3800 system equipped with capillary GC column of DB-5 (5% diphenyl, 95% dimethyl polysiloxane) type with 30 m length, 0.25 mm diameter and 0.25 *μ*m film thickness. The injection volume was 1 *μ*L. The temperature was programmed as follows: 60° C for 1 min and then increased gradually up to 246° C at 3 °C/min until the end of the run of 66 min. Ultrapure helium gas (99.999%) was used as a carrier gas at a flow rate of 1 mL/min; split ratio of 1:40. Scanning range for mass detector was from 35 to 500 *m/z*. Analysis was done in duplicates. The volatile compounds were identified using both their mass spectra and retention indices, arithmetic Kovat’s index (AKI), calculated relative to a hydrocarbon mixture of n-alkane (C_8_-C_20_) analyzed under the same chromatographic conditions of sample analysis. For every compound separated in the GC-MS and by using its retention time as well as the retention times of the n-alkane (C_8_-C_20_) hydrocarbon mixture, the AKI was calculated using the Van Den Dool and Kratz equation [38]. A computer program was used for matching compounds’ mass spectra (MS) with NIST, ADAMS, and WILEY libraries. The identification was further confirmed by matching the calculated arithmetic indices with those reported in the literature [27]. The percentage composition of the individual components was calculated by a normalization process, utilizing their relative peak areas, assuming a unity response by all components.

#### 3.4 Hierarchical cluster analysis

The published chemical composition of the essential oil of *H. triquetrifolium* grown wildly in several geographical locations in the world were analyzed using Hierarchical Cluster Analysis (HCA). After a thorough literature review, the tabulated essential oil contents with their corresponding concentrations in each report were pooled into one table, which consisted of 312 chemical compounds. Each reported essential oil sample was considered as an independent operational taxonomic unit. The concentration of non-reported chemical compounds was considered zero. Chemical compounds found in traces were assigned a concentration of 0.01%. Agglomerative hierarchical clustering was performed using XLSTAT software v.2014 (Addinsoft, New York, USA) as previously described [39, 40]. Pearson’s correlation was utilized for the similarity assignment and clusters were defined by the unweighted pair-group method.

### 3.5 Antibacterial activity of H. triquetrifolium essential oil

*H. triquetrifolium* essential oil was evaluated for antibacterial activity using the disc diffusion [41] and micro broth dilution [42] assays. In disk diffusion assay, the density of the bacterial suspension was adjusted to 1.0 × 10^8^ UFC/mL (or 0.5 McFarland turbidity units). A sterile cotton swab was used to inoculate the entire surface of the Mueller-Hinton agar plate. The following bacterial strains were used for testing: *Escherichia coli, Staphylococcus aureus, Staphylococcus chromogenes*, and *Bacillus subtilis*. All strains were isolated from cows’ milk that were affected with clinical mastitis. The essential oil, dissolved in DMSO (1:1), was applied on a sterile disc and then aseptically placed on the inoculated plates. The plates were left at room temperature for 1 h and then incubated at 37 °C for 18-24 h. Amoxicillin (10 μg/disc) and enrofolxacin (5 μg/disc) were used as positive controls, and DMSO as a negative control. After the 24 h incubation period, the zones of inhibition were measured in millimeters. All experiments were performed in duplicates.

The minimal inhibitory concentrations (MICs) and the minimum bactericidal concentrations (MBC) were determined by micro broth dilution assay [42] using a 96-well plate. Multiple bacterial suspensions of *E. coli, S. aureus, S. chromogenes*, and *B. subtilis* were prepared at a concentration of 1.0 × 10^8^ UFC/mL. A stock solution of *H. triquetrifolium* essential oil (80%) was prepared by dissolving in DMSO. Different DMSO concentrations were tested to avoid any antibacterial effect that could be attributed to DMSO. The final concentration of DMSO was never exceeded 1%. About 100 *μ*L of broth was placed in each well, and then 100 *μ*L of the oil stock solution was added to the first well. Different concentrations of the essential oil stock were obtained using distilled water as follows: 100%, 50%, 25%, 12.5%, 6.2%, and 3.1% (v/v).

The plate was left at room temperature for 1 h, followed by incubation 37 °C for 18-24 h. Turbidity of wells is used as an indicator to determine MIC and MBC. MIC is defined as the first well with no visible bacterial growth, while MBC is the first well that showed no growth on solid media after 24 h incubation. In our experiments, turbidity was not conclusive and somehow confusing. Hence, about 50 *μ*L from each well was transferred into agar plates and incubated at 37 °C for 18-24 h. MBC was considered as the lowest concentration of the oil that allowed less than 10 colonies to grow in the agar plate.

## 4. Conclusion

The current study has identified the chemical composition of the essential oil extracted from the aerial parts of *H. triquetrifolium* for the first time in Jordan. It shows that the chemical profile is dominated by monoterpene hydrocarbons. The principal monoterpene hydrocarbon identified was *α*-pinene, which could be identified as a chemotype of Jordanian *H. triquetrifolium*. Hierarchical Cluster Analysis of the essential oil composition of *H. triquetrifolium* from different geographical locations in the world has identified the oil from Jordan to be more similar to those obtained from Italy, Greece, and Crete. The antimicrobial evaluation of the extracted oil indicates a potent antimicrobial activity against *E. coli, S. aureus, S. chromogenes*, and *B. subtilis*. Further studies are essential to address the safety of *H. triquetrifolium* essential oil along with other pharmacological uses.

## References

[1] O.G. Roskov Y., Orrell T., Nicolson D., Bailly N., Kirk P.M., Bourgoin T., DeWalt R.E., Decock W., Nieukerken E. van, Zarucchi J., Penev L., eds. (2019). Species 2000 & ITIS Catalogue of Life, 2019 Annual Checklist. Digital resource at https://www.catalogueoflife.org/annual-checklist/2019. Species 2000: Naturalis, Leiden, the Netherlands. ISSN 2405-884X. Published 2019.

[2] N.K.B. Robson, in: Flora Europaea, edited by T.G. Tutin, V.H. Heywood, N.A. Burges, D.M. Moore, D.H. Valentine, S.M. Walters and D.A. Webb (Cambridge University Press, Cambridge, 1968).

[3] M.Á. Alonso, J.C. Agulló, J.L. Villar, A. Juan, and M.B. Crespo. presented at: Annales Botanici Fennici; 2013.

[4] J.M. Greeson, B. Sanford, and D.A. Monti. St. John’s wort (Hypericum perforatum): a review of the current pharmacological, toxicological, and clinical literature. Psychopharmacology. 153, 402 (2001).

[5] D. Al-Eisawi, Flora of Jordan Checklist, 1st ed. (The University of Jordan Press, Amman, Jordan, 2013).

[6] W. Dhifi, S. Bellili, S. Jazi, N. Bahloul, and W. Mnif. Essential oils’ chemical characterization and investigation of some biological activities: A critical review. Medicines (Basel). 3, 25 (2016).

[7] P.F. Stevens, in: The Families and Genera of Vascular Plants, edited by K. Kubitzki (Springer, Berlin, Heidelberg, 2007).

[8] F.Q. Alali and K. Tawaha. Dereplication of bioactive constituents of the genus hypericum using LC-(+,−)-ESI-MS and LC-PDA techniques: Hypericum triquterifolium as a case study. Saudi Pharm J. 17, 269 (2009).

[9] K. Tawaha, F.Q. Alali, M. Gharaibeh, M. Mohammad, and T. El-Elimat. Antioxidant activity and total phenolic content of selected Jordanian plant species. Food Chem. 104, 1372 (2007).

[10] L. Pistelli, A. Bertoli, I. Morelli, F. Menichini, R.A. Musmanno, T.D. Maggio, and G. Coratza. Chemical and antibacterial evaluation of Hypericum triquetrifolium Turra. Phytother Res. 19, 787 (2005).

[11] B. Ozturk, S. Apaydin, E. Goldeli, I. Ince, and U. Zeybek. Hypericum triquetrifolium Turra. extract exhibits antiinflammatory activity in the rat. J Ethnopharmacol. 80, 207 (2002).

[12] F. Conforti, M.R. Loizzo, A.G. Statti, and F. Menichini. Cytotoxic activity of antioxidant constituents from Hypericum triquetrifolium Turra. Nat Prod Res. 21, 42 (2007).

[13] M. Couladis, P. Baziou, E. Verykokidou, and A. Loukis. Antioxidant activity of polyphenols from Hypericum triquetrifolium Turra. Phytother Res. 16, 769 (2002).

[14] F. Conforti, G.A. Statti, R. Tundis, F. Menichini, and P. Houghton. Antioxidant activity of methanolic extract of Hypericum triquetrifolium Turra aerial part. Fitoterapia. 73, 479 (2002).

[15] S. Apaydın, U. Zeybek, I. Ince, G. Elgin, C. Karamenderes, B. Ozturk, and I. Tuglular. Hypericum triquetrifolium Turra. extract exhibits antinociceptive activity in the mouse. J Ethnopharmacol. 67, 307 (1999).

[16] F. Alali, K. Tawaha, and T. Al-Eleimat. Determination of hypericin content in Hypericum triquetrifolium Turra (Hypericaceae) growing wild in Jordan. Nat Prod Res. 18, 147 (2004).

[17] K. Hosni, K. Msaada, M. Ben Taârit, and B. Marzouk. Phenological variations of secondary metabolites from Hypericum triquetrifolium Turra. Biochem Syst Ecol. 39, 43 (2011).

[18] F. Karim and S. Quraan, Medicinal Plants of Jordan. (Yarmouk University, Irbid, Jordan, 1986).

[19] S.L. Crockett. Essential oil and volatile components of the genus Hypericum (Hypericaceae). Nat Prod Commun. 5, 1493 (2010).

[20] H. Karim, M. Kamel, B.T. Mouna, C. Thouraya, and M. Brahim. Essential oil composition of Hypericum triquetrifolium Turra. aerial parts. Ital J Biochem. 56, 40 (2007).

[21] G. Kızıl, Z. Toker, H.Ç. Özen, and Ç. Aytekin. The antimicrobial activity of essential oils of Hypericum scabrum, Hypericum scabroides and Hypericum triquetrifolium. Phytother Res. 18, 339 (2004).

[22] Z. Rouis, N. Abid, S. Koudja, T. Yangui, A. Elaissi, P.L. Cioni, G. Flamini, and M. Aouni. Evaluation of the cytotoxic effect and antibacterial, antifungal, and antiviral activities of Hypericum triquetrifolium Turra essential oils from Tunisia. BMC Complementary Altern Med. 13, 24 (2013).

[23] Z. Rouis, A. Elaissi, N.B.S. Abid, M.A. Lassoued, P.L. Cioni, G. Flamini, and M. Aouni. Chemical composition and intraspecific variability of the essential oils of five populations of Hypericum triquetrifoliumTurra Growing in North Tunisia. Chem Biodiversity. 9, 806 (2012).

[24] S.E. Sajjadi, I. Mehregan, and M. Taheri. Essential oil composition of Hypericum triquetrifolium Turra growing wild in Iran. Res Pharm Sci. 10, 90 (2015).

[25] E. Yuce and E. Bagci. The essential oils of the aerial parts of two Hypericum taxa (Hypericum triquetrifolium and Hypericum aviculariifolium subsp. depilatum var. depilatum (Clusiaceae)) from Turkey. Nat Prod Res. 26, 1985 (2012).

[26] D. Al-Eisawi, Field Guide to Wild Flowers of Jordan and Neighbouring Countries, 1st ed. (Jordan Press Foundation, Amman, Jordan, 1998), p. 296.

[27] R.P. Adams, Identification of Essential Oil Components By Gas Chromatography/Mass Spectrometry, 4th ed. (Allured Pub Corp, Carol Stream, IL, 2007).

[28] P.V. Petrakis, M. Couladis, and V. Roussis. A method for detecting the biosystematic significance of the essential oil composition: The case of five Hellenic Hypericum L. species. Biochem Syst Ecol. 33, 873 (2005).

[29] A. Bertoli, F. Menichini, M. Mazzetti, G. Spinelli, and I. Morelli. Volatile constituents of the leaves and flowers of Hypericum triquetrifolium Turra. Flavour Fragrance J. 18, 91 (2003).

[30] K. Hosni, K. Msaada, T. Chahed, M. Ben Taarit, and B. Marzouk. Comparative analysis of the essential oils of Hypericum triquetrifolium Turra. Extracted by ultrasound, hydrodistillation and soxhlet/dynamic headspace. Med Aromat Plant Sci Biotechnol. 5, 100 (2010).

[31] Z. Rouis, A. Elaissi, N.B.S. Abid, M.A. Lassoued, P.L. Cioni, G. Flamini, and M. Aouni. Chemical Composition and Intraspecific Variability of the Essential Oils of Five Populations of Hypericum triquetrifolium Turra Growing in North Tunisia. Chemistry & Biodiversity. 9, 806 (2012).

[32] Z. Rouis, N. Abid, S. Koudja, T. Yangui, A. Elaissi, P.L. Cioni, G. Flamini, and M. Aouni. Evaluation of the cytotoxic effect and antibacterial, antifungal, and antiviral activities of Hypericum triquetrifolium Turra essential oils from Tunisia. BMC Complementary and Alternative Medicine. 13, 24 (2013).

[33] S.E. Sajjadi, I. Mehregan, and M. Taheri. Essential oil composition of Hypericum triquetrifolium Turra growing wild in Iran. Res Pharm Sci. 10, 90 (2015).

[34] H. Azeez. Comparison between Hypericum triquetrifolium leaves and derived calli in essential oil content. Journal of Al-Nahrain University-Science. 20, 123 (2017).

[35] K. Hosni, K. Msaada, M. Ben Taarit, T. Chahed, and M. Brahim. Essential oil composition of Hypericum triquetrifolium Turra. aerial parts. Ital J Biochem. 56, 40 (2007).

[36] A.C. da Silva Rivas, P.M. Lopes, M.M. de Azevedo Barros, D.C. Costa Machado, C.S. Alviano, and D.S. Alviano. Biological activities of α-pinene and β-pinene enantiomers. Molecules. 17, 6305 (2012).

[37] British Pharmacopoeia Commission, British Pharmacopoeia. (London, HMSO, 1988), p. 137–138.

[38] H. van Den Dool and P. Dec. Kratz. A generalization of the retention index system including linear temperature programmed gas—liquid partition chromatography. J Chromatogr A. 11, 463 (1963).

[39] P. Satyal, D.J. Craft, S.N. Dosoky, and N.W. Setzer. The chemical compositions of the volatile oils of garlic (Allium sativum) and wild garlic (Allium vineale). Foods. 6, (2017).

[40] A. Assaeed, A. Elshamy, E.A. El Gendy, B. Dar, S. Al-Rowaily, and A. Abd-ElGawad. Sesquiterpenes-rich essential oil from above ground parts of Pulicaria somalensis exhibited antioxidant activity and allelopathic effect on weeds. Agronomy. 10, (2020).

[41] Clinical and Laboratory Standards Institute (CLSI), Performance Standards for Antimicrobial Disk Susceptibility Tests, 10th ed. (Clinical and Laboratory Standards Institute, Wayne, PA, 2018), p. 15–39.

[42] linical and Laboratory Standards Institute (CLSI), Method for Dilution Antimicrobial Susceptibility Tests for Bacteria that Grow Aerobically, 11th ed. (Clinical and Laboratory Standards Institute, Wayne, PA, 2018), p. 15–50.

